# Clusterin reverses epitheliopathy, reduces inflammation, and restores goblet cells and corneal nerves in a mouse model of autoimmune dry eye

**DOI:** 10.1101/2025.06.05.658110

**Authors:** Jonas Franz, Tat Fong Ng, Shivali Gupta, Mark R. Wilson, M. Elizabeth Fini, Sharmila Masli

## Abstract

Chronic ocular surface disease (OSD) is characterized by corneal epitheliopathy, reduced barrier function and loss of nerves, accompanied by persistent inflammation. Current treatments offer limited relief and there is no approved therapy that promotes neurosensory regeneration in OSD. Here, we tested the therapeutic efficacy of clusterin (CLU), a molecular chaperone and MMP9 inhibitor found in tears, in *Thbs1*-deficient mice, a preclinical model of autoimmune dry eye associated with Sjögren’s disease (SjD). These mice were treated topically at the ocular surface, bilaterally, for 3 weeks with recombinant human CLU (rhCLU) or human plasma-derived CLU (pCLU) eyedrops and compared to standard-of-care 0.1% dexamethasone eyedrops. Treatment with CLU significantly improved corneal barrier integrity, increased corneal nerve density, enhanced the proportion of corneal nerves with immunoreactivity for CGRP and promoted conjunctival goblet cell regeneration. Furthermore, CLU reduced immunoreactivity for ADAM17 in the corneal epithelium and reduced *Tnfa* expression in the conjunctiva, supporting its anti-inflammatory effect. Notably, all these effects were comparable to, or even exceeded, those resulting from treatment with dexamethasone. Based on its efficacy, we introduce CLU as a multifunctional and promising biotherapeutic for a widespread range of ocular inflammatory conditions involving corneal epitheliopathy and nerve loss, including dry eye associated with SjD.

## Introduction

Sjögren’s disease (SjD) is a chronic autoimmune disorder characterized by inflammatory destruction and functional loss of the lacrimal glands, leading to reduced tear production and subsequent dry eye disease ^1–3^. This tear deficiency results in ocular surface inflammation, including epithelial defects in both the cornea and conjunctiva, a condition clinically termed as keratoconjunctivitis sicca ^4^. Patients often experience burning, stinging and itching sensations in the eyes, along with blurred vision. ^4–6^. These symptoms correlate with the altered corneal nerve morphology and reduced nerve density observed in SjD patients ^7–10^. Managing ocular symptoms in these patients remain a significant challenge. Currently approved treatments for dry eye include artificial tears, which provide only temporary symptomatic relief, and anti-inflammatory drugs, which often yield suboptimal results and are associated with significant limitations, including notable side effects ^2,11,12^. Importantly, no approved treatment specifically addresses the neurosensory damage of the ocular surface that contributes to dry eye symptoms. Therefore, finding new, targeted and specific therapeutic alternatives to treat neurosensory damage in dry eye is of great clinical interest.

The cornea plays a critical role in maintaining ocular health, serving as a transparent, densely innervated outer layer that protects internal structures and contributes significantly to the eye’s refractive power ^13^. Its biological functions rely on the tight cooperation among cellular components, including the corneal epithelium and sensory nerve fibers ^14,15^. Neurotransmitters such as CGRP, released by corneal nerves, promote corneal epithelial proliferation and healing ^16,17^, while corneal epithelial cells, in turn, support nerve regeneration by producing neurotrophic factors like NGF and GDNF ^18,19^. This bidirectional trophism ensures effective recovery and regeneration of both the nerves and the epithelium, preserving the protective barrier and sensory functions of the eye. Such a reciprocal relationship is especially crucial during wound healing when the epithelial layer is damaged ^19^. Dry eye disease, trauma, infections or decreased tear production can lead to those disruptions in the epithelial barrier, also known as ocular epitheliopathy ^6,20^.

CLU is an evolutionarily conserved, homeostatic, secreted glycoprotein expressed by mucosal epithelia at fluid-tissue interfaces and found in all bodily fluids ^21–24^. At the ocular surface, *CLU* is expressed by epithelial cells and in the lacrimal gland and CLU protein can be detected in tears ^21,22,25–28^. CLU is cytoprotective, anti-inflammatory and proteostatic ^21–23^, acting as both a molecular chaperone ^29,30^ and an inhibitor of MMP9 ^31^, a critical mediator of proteolytic damage to the ocular surface epithelia in dry eye disease ^32–34^. In acutely damaged corneas, CLU selectively binds to the damaged ocular surface, sealing the epithelial barrier and preventing further damage to the cells ^35–37^. In humans and mice with aqueous-deficient dry eye disease, CLU levels in the tears decrease, rendering the ocular surface more vulnerable to damage ^35,38^. However, it remains unknown whether CLU is anti-inflammatory in the context of dry eye, or whether it might contribute to the repair and recovery of damaged corneal nerves lost to chronic inflammation in the disease.

Thrombospondin-1 (*Thbs1*)-deficient mice have become an established and valuable preclinical model for studying SjD dry eye disease ^39^. These mice spontaneously develop SjD-associated chronic ocular surface inflammation like that observed in SjD patients ^39^, including corneal epitheliopathy and the loss of mucin-secreting goblet cells ^39–42^. Concurrently, a significant reduction in corneal nerve density is observed in *Thbs1*-deficient mice, mirroring findings in SjD patients ^10,43^. This corneal nerve abnormality is also marked by significantly reduced expression of neurotransmitters like CGRP, which is known to protect corneal epithelial cells through its cytoprotective, pro-regenerative and anti-inflammatory roles ^16,17^. Overall, the clinical and pathological similarities between *Thbs1*-deficient mice and SjD patients make these mice a powerful preclinical model for investigating the efficacy of CLU in repairing corneal epithelial and nerve damage.

In this study, we investigated the therapeutic efficacy of topically applied CLU in *Thbs1*-deficient mice with established ocular surface disease. We used human CLU produced in our laboratory by recombinant DNA methodologies (rhCLU) at two different doses, comparing it to human plasma-derived CLU (pCLU) and to 0.1% dexamethasone, a standard of care.

## Results

### Topical application of clusterin improves corneal epithelial barrier in a preclinical mouse model of Sjögreńs disease

To assess the efficacy of CLU in repairing corneal epithelial damage associated with SjD, we utilized *Thbs1*-deficient mice, a well-established preclinical model of SjD-related ocular surface disease ^39–41^. In this model, corneal epithelial damage becomes fully evident by 12 weeks of age. Therefore, we conducted our double-blinded preclinical study using 12-week-old *Thbs1*-deficient mice. Different groups of male and female *Thbs1*-deficient mice were topically treated twice a day, five days per week (skipping weekends), for three weeks with either rhCLU (1 µg/ml or 50 µg/ml) made in our laboratory, pCLU (50 µg/ml), or vehicle control (PBS). We compared these treatments to treatment with dexamethasone 0.1%, the current standard of care ^44^, which served as a positive control due to its well-documented anti-inflammatory effects. Corneal epithelial damage was assessed before treatment initiation (baseline), once per week, and at the study endpoint using corneal fluorescein staining (CFS) (**Figure 1A**). Normalized CFS scores were compared across different treatment groups (**Figure 1B, 1C**). In vehicle-treated mice, CFS scores progressively increased each week, consistent with the age-related disease progression as previously reported in this model ^41,45^. Conversely, in mice treated with 1 µg/ml rhCLU, the weekly CFS scores remained largely stable, showing neither progression nor improvement (**Figure 1B**). However, at higher concentration of 50 µg/ml, both rhCLU and pCLU treatments led to a significant epithelial barrier improvement, as evidenced by a significant decrease in CFS. This improvement was already detectable after one week of treatment and continued to progress until the end of the study. A similar pattern of corneal epithelial improvement was observed in both male and female mice treated with 50 µg/ml of either rhCLU or pCLU (**Figure 1B, 1C**). When comparing overall treatment efficacy in repairing corneal epithelial damage, both rhCLU and pCLU demonstrated improvements comparable to those achieved with 0.1% dexamethasone, irrespective of sex (**Figure 1D, 1E**). Finally, we examined whether CLU treatment induced any toxic morphological changes or inflammatory cell infiltrations in the cornea and conjunctiva. Hematoxylin and Eosin (H&E) staining revealed no morphological alterations or inflammatory infiltrates in these tissues following CLU application (**Supplementary Figure 1**).

**Figure 1.**
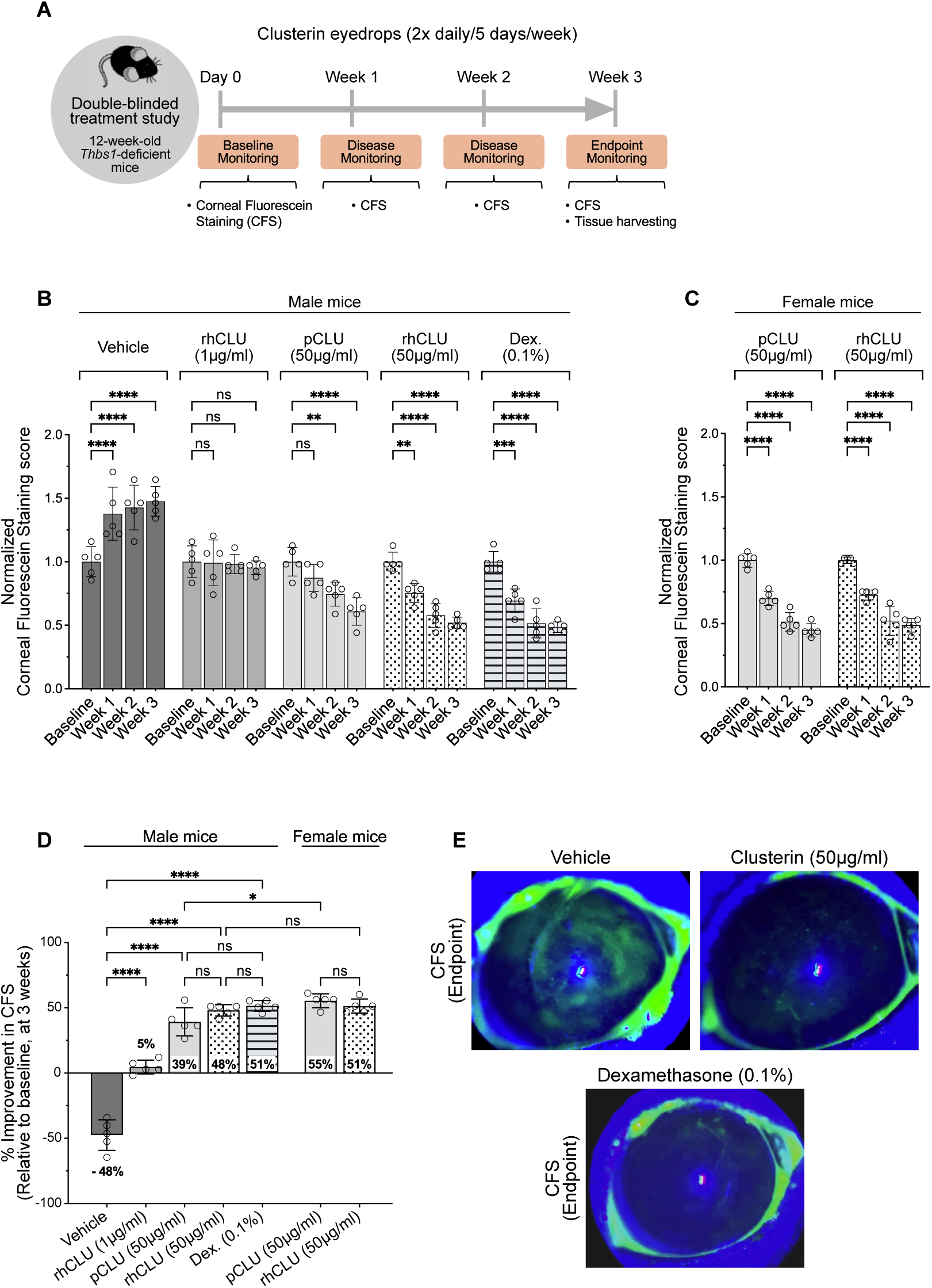
Topical application of CLU improves corneal epithelial barrier in a preclinical mouse model of Sjögreńs disease. (**A**) Scheme illustrates timeline of the double-blinded study. 12 weeks-old Thrombospondin-1 (*Thbs1*) deficient mice were used as a preclinical model of chronic ocular surface inflammation associated with Sjögreńs disease. Mice (n= 5/group) were treated bilaterally, b.i.d. with vehicle, rhCLU (1 µg/ml or 50 µg/ml), pCLU (50 µg/ml), or 0.1% dexamethasone (positive control) for 3 weeks. Disease progression was monitored before starting (baseline) and after 1, 2 and 3 weeks of treatment by assessing corneal epitheliopathy with corneal fluorescein staining (CFS). (**B, C**) Normalized weekly CFS scores of male and female *Thbs1*-deficient mice with indicated treatments. (**D**) Relative improvement in CFS with each treatment. (**E**) Representative images of CFS (green staining) taken at the endpoint of the study. Error bars represent ± SEM (ns ≥ 0.05, *p < 0.05, **p < 0.01, ***p < 0.001, ****p < 0.0001).

Taken together, these results demonstrate that the previously reported corneal epithelial sealing properties of CLU effectively improve corneal barrier integrity, even under chronic inflammatory conditions, in *Thbs1*-deficient mice.

### Corneal nerve density is reduced in *Thbs1*-deficient mice, as seen in SjD patients

Significant reduction in corneal nerve density has been previously observed in SjD patients ^8,10^. A similar change was also reported in 12-week-old *Thbs1*-deficient mice ^46^. Here, we confirmed reduced corneal nerve density in 15-wk-old *Thbs1*-deficient mice, which matched the age of the mice used in our study at the endpoint. Corneas from 15-week-old *Thbs1*-deficient and wild-type (WT) mice were harvested and immunostained for beta-3-tubulin, a component of the neuronal tubulin cytoskeleton ^47,48^, as well as for the neurotransmitter CGRP. Central and peripheral corneas have different patterns of subbasal nerve organization ^47^. Therefore, in this study, to gain a deeper understanding of nerve morphology changes, we analyzed the central and peripheral corneas separately (**Figure 2A**). As shown in **Figure 2B**, the localization of subbasal nerves in the central cornea exhibited a typical densely-packed, vortex-like pattern ^47,49^. Quantitative assessment and comparison of beta-3-tubulin-stained nerve densities between WT and *Thbs1*-deficient mice revealed a significant reduction in nerve density in *Thbs1*-deficient mice (**Figure 2C**). In contrast, in the periphery, corneal nerves predominantly ran parallel to each other (**Figure 2D**), forming a complex, well-organized network extending to the limbal area. Both nerve density and organization were significantly disrupted in peripheral corneas of *Thbs1*-deficient mice, as evidenced by large nerve-free gaps in the whole mount images and the loss of fine nerve structures (**Figure 2D**). Quantitative analysis of the nerve density, both in the central and peripheral regions, confirmed a significant reduction in *Thbs1*-deficient mice compared to WT (**Figure 2E**). Interestingly, the decline in peripheral corneal nerve density in *Thbs1*-deficient mice was nearly five times greater than that observed in the central cornea. In addition, the relative level of immunoreactive CGRP was notably reduced in both the central and peripheral corneas of *Thbs1*-deficient mice compared to WT mice **(Figure 2F-2H**). These results further confirm corneal nerve abnormalities in *Thbs1*-deficient mice ^46^, mirroring the findings in SjD patients and supporting the use of this mouse model to evaluate effects of topically delivered CLU on corneal nerve alterations and related biology.

**Figure 2.**
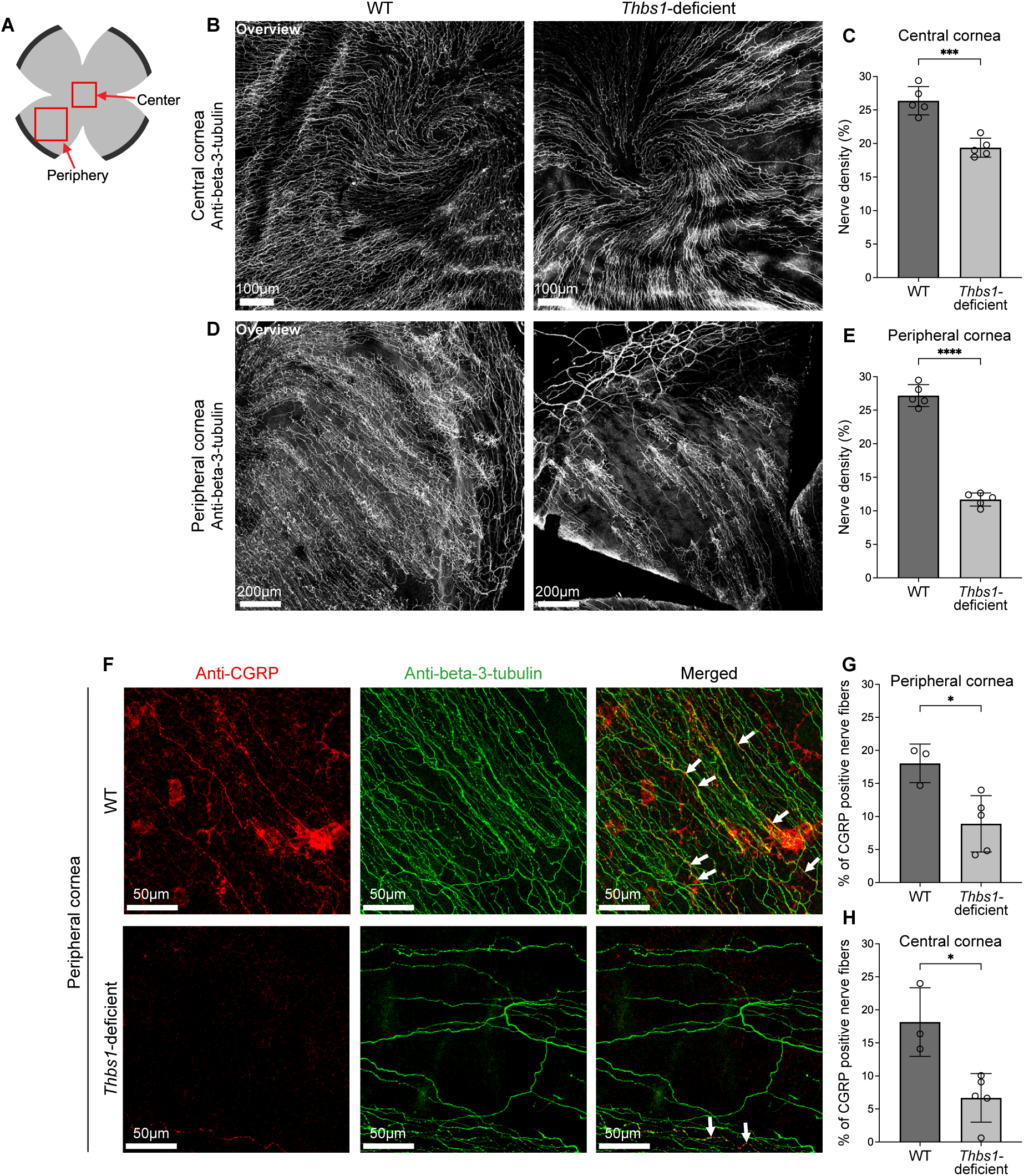
Nerve density and proportion of CGRP-positive nerves are reduced in the central and peripheral corneas of mice with Sjögreńs disease. Corneal nerve density and CGRP-positive nerves in peripheral and central cornea of WT and *Thbs1*-deficient mice were evaluated by confocal microscopy in wholemount corneas immunostained with antibodies against pan-neuronal marker, beta-3-tubulin, and neurotransmitter CGRP. (**A**) Z scans from peripheral and central area of flattened wholemount corneas (WT n= 5 and *Thbs1*-deficient n= 5) were analyzed; (**B, D**) Representative confocal images of the overview of staining for beta-3-tubulin in (**B**) the central and (**D**) peripheral cornea of WT and *Thbs1*-deficient mice and (**C, E**) corresponding quantitative analysis of the images. Nerve density represents the percentage of the total image area occupied by positively stained nerves. (**F**) Representative confocal images from the peripheral cornea of WT and *Thbs1*-deficient mice immunostained for CGRP and beta-3-tubulin. (**G, H**) Corresponding quantitative analysis of CGRP-expressing nerves in WT (n= 3) and *Thbs1*-deficient mice (n= 5). Percentage of CGRP positive nerves was determined based on the ratio of CGRP positive nerve length to total beta-3-tubulin positive nerve length in each image. Arrows in merged images indicate CGRP positive nerve fibers. Error bars indicate ± SEM (*p <0.05, ***p < 0.001, ****p < 0.0001).

### Topically applied CLU improves corneal nerve density in mice with Sjögreńs disease

To evaluate whether CLU can also mediate corneal nerve regeneration, we analyzed corneal nerve densities in *Thbs1*-deficient mice after the different treatment approaches, as outlined earlier. At the study endpoint, corneas from groups treated with vehicle, 1 µg/ml of rhCLU, 50 µg/ml of rhCLU, pCLU or 0.1% dexamethasone were harvested and immunostained for beta-3-tubulin and CGRP. Quantitative analysis was then performed on beta-3-tubulin-stained corneal nerves, as well as the proportion of CGRP-positive nerves within the corneal nerve population. Representative immunofluorescence images of stained corneas indicated that, compared to vehicle-treated mice, those treated with rhCLU (50 µg/ml) or 0.1% dexamethasone displayed improvements in nerve densities in both the peripheral and central corneas (**Figure 3A, 3B**). Indeed, quantitative analyses confirmed that nerve densities in the peripheral cornea of mice treated with 50 µg/ml of either rhCLU or pCLU - but not 1 µg/ml of rhCLU - were significantly improved compared to vehicle-treated mice (**Figure 3C**). Treatment with 0.1% dexamethasone also significantly improved corneal nerve density. Strikingly however, treatment with either rhCLU or pCLU at 50 µg/ml resulted in a significantly greater improvement in peripheral corneal nerve density than dexamethasone (**Figure 3C**). In contrast, in the central cornea, only pCLU (50 µg/ml) treatment led to a significant improvement in nerve density compared to either vehicle or dexamethasone treatment (**Figure 3D**). Although treatment with 50 µg/ml of rhCLU showed a trend toward improved central corneal nerve density, quantitative analysis did not reveal a statistically significant change (**Figure 3D**). Taken together, in comparison to the vehicle-treated group, nerve density improvements were notably more prominent in the peripheral cornea than in the central cornea, while the most improvement was achieved by CLU treatment.

**Figure 3.**
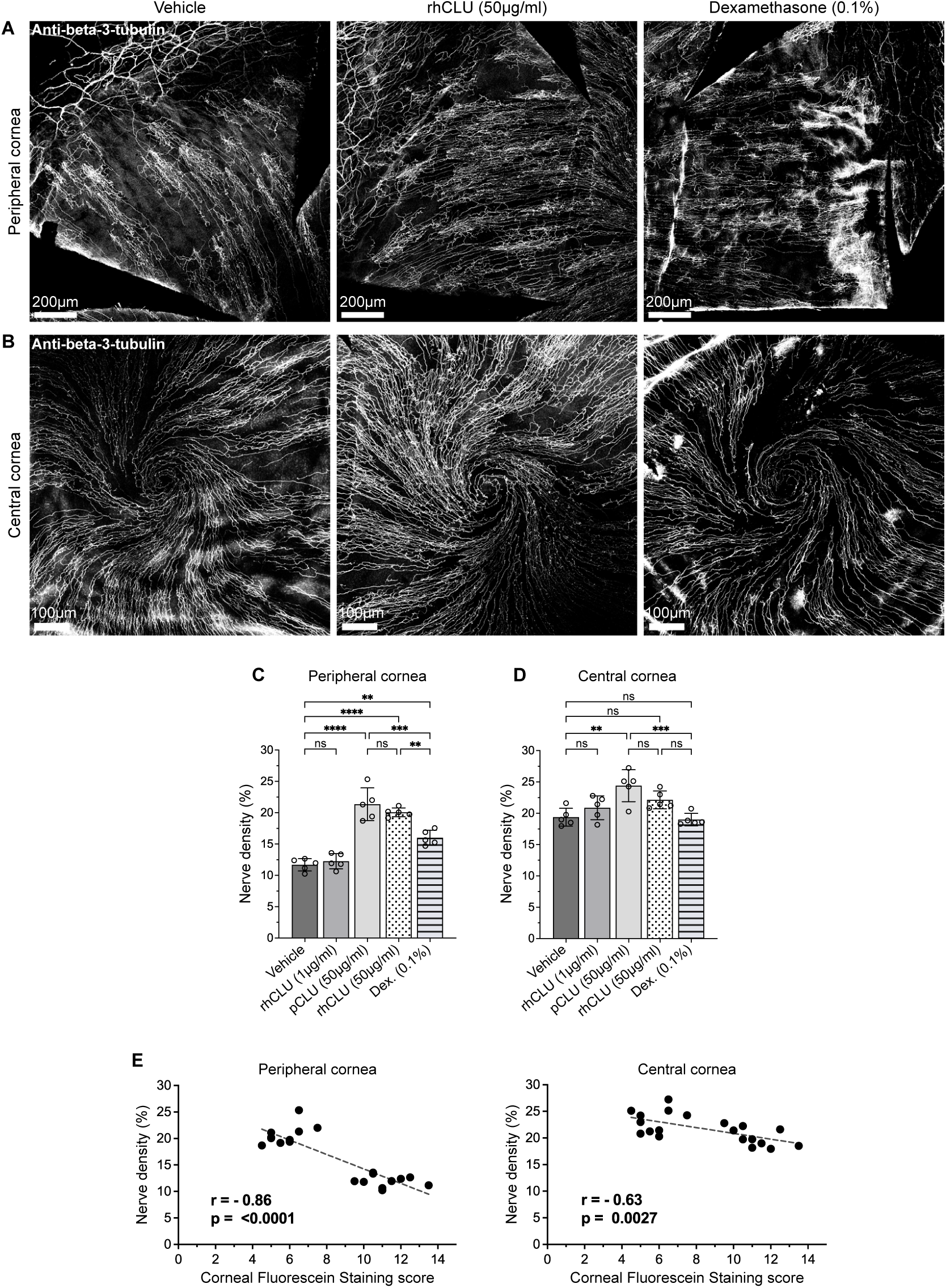
Topically applied CLU improves corneal nerve density in mice with Sjögreńs disease. Corneas harvested at the study end point (3 weeks) were immunostained with anti-beta-3-tubulin. (**A, B**) Representative overview confocal images from indicated treatment groups and corresponding quantitative analysis of (**C**) peripheral and (**D**) central corneal nerves. Nerve density was analyzed as the percentage of beta-3-tubulin positive nerves relative to the total image area (n= 5 mice/group). (**E**) Corneal epithelial barrier improvement correlates with the enhancement in corneal nerve density. Corneal Fluorescein Staining score (CFS) for each mouse was plotted against the corresponding corneal nerve density in central and peripheral cornea (n= 20 mice). Spearman correlation (r) analysis revealed a significant negative correlation between CFS and nerve density. Error bars represent ± SEM (ns ≥ 0.05, **p < 0.01, ***p < 0.001, ****p < 0.0001).

Additionally, we also assessed a potential correlation between corneal epithelial integrity, as reflected by CFS scores, and corneal nerve density. As shown in **Figure 3E**, a significant negative correlation was found between CFS scores and corneal nerve density in both the peripheral and central corneal regions. Specifically, lower CFS scores (indicating improved epithelial barrier) are associated with higher nerve density in mice with SjD. This correlation is consistent with the known reciprocal trophism between the corneal epithelium and subbasal nerves in patients with SjD neurotrophic keratopathy ^10,50^.

### Topical CLU increases the proportion of CGRP-positive corneal nerves

A reduced proportion of CGRP-positive corneal nerves in *Thbs1*-deficient mice is consistent with the reported wound-healing and anti-inflammatory properties of this neuropeptide ^17,51^. To determine if CLU-driven repair of corneal epithelial damage also helps restore CGRP-positive corneal nerves, we performed further quantification of immunoreactive CGRP in beta-3-tubulin-stained corneal nerves. This analysis revealed an increased proportion of beta-3-tubulin-stained nerves that were also stained positively for CGRP in mice treated with rhCLU (50 µg/ml) or 0.1% dexamethasone, as indicated by confocal imaging (**Figure 4A**) and the corresponding quantitative data. Enhanced CGRP positivity was observed in both the peripheral and central regions of the cornea (**Figure 4B, 4C**). In conclusion, our results demonstrate that, in addition to repairing corneal epithelial damage and improving corneal nerve density, CLU also enhances the proportion of neuropeptide CGRP-positive corneal nerves.

**Figure 4.**
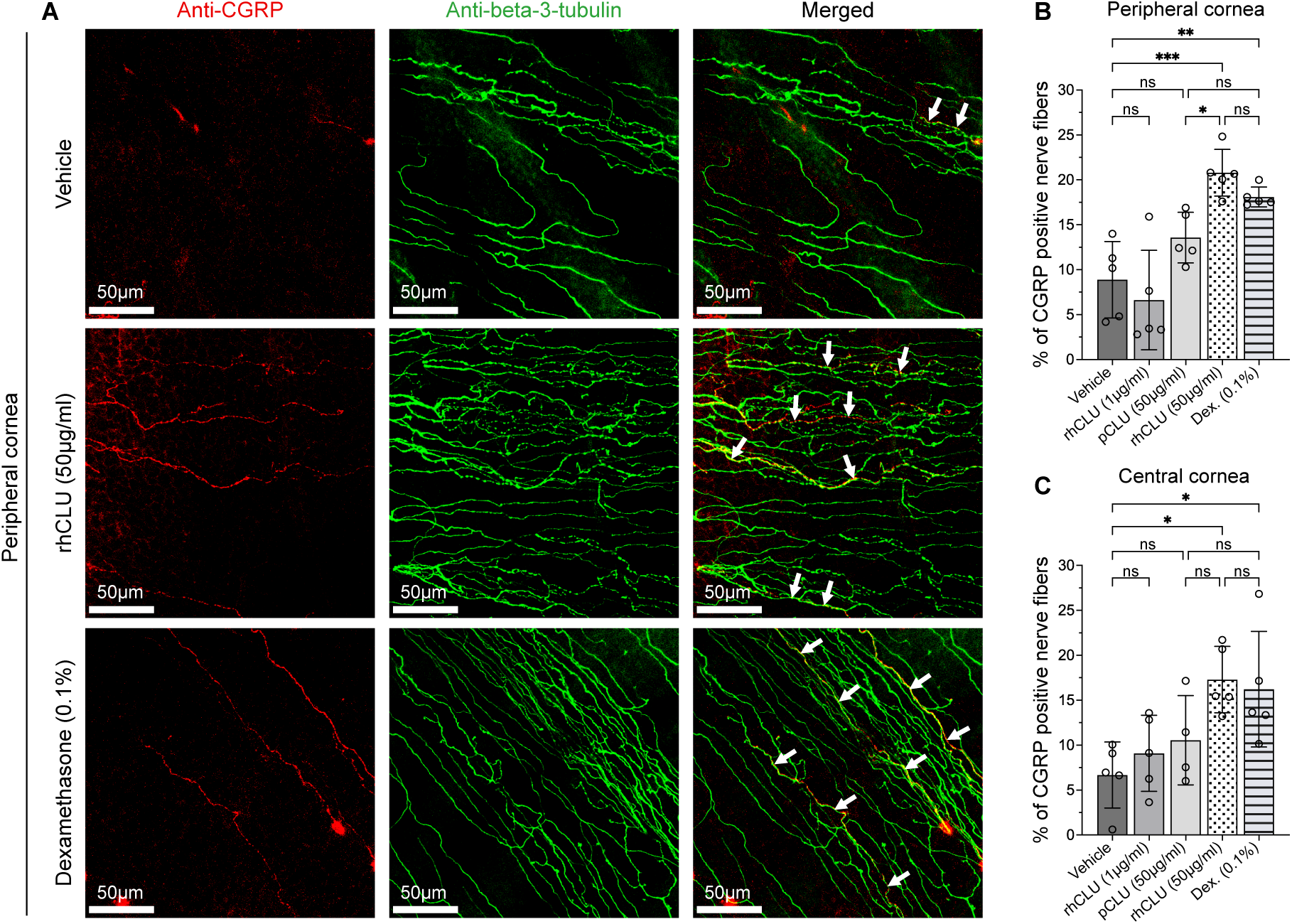
Topical CLU increases the proportion of CGRP-positive corneal nerves. (**A**) Representative immunostainings showing anti-CGRP and anti-beta-3-tubulin in peripheral cornea from mice with indicated treatments at the study endpoint (3 weeks). (**B, C**) Quantitative analysis of CGRP-expressing nerves in the (**B**) peripheral and (**C**) central corneas (n= 5 mice/group). Arrows in merged images indicate CGRP positive nerve fibers. Error bars represent ± SEM (ns ≥ 0.05, *p < 0.05, **p < 0.01, ***p < 0.001).

### CLU treatment helps counter SjD-associated conjunctival inflammation

As reported in humans, chronic dry eye associated with SjD in *Thbs1*-deficient mice is characterized by a substantial loss of conjunctival goblet cells, driven by local inflammation ^41,52,53^. To determine whether topically applied CLU also helps mitigate conjunctival inflammation and aids in the recovery of goblet cell density, mucin-filled goblet cells stained with Alcian Blue in histological sections of the conjunctiva were enumerated. Tissues harvested at the study endpoint from the various treatment groups were examined. Overall, increased numbers of goblet cells were detectable in mice treated with rhCLU, pCLU or dexamethasone, as compared to vehicle-treated control mice (**Figure 5A**). Quantitative analysis of goblet cells confirmed significantly increased numbers in mice treated with 0.1% dexamethasone (Figure 5B), as reported previously ^41^. In addition, treatment with rhCLU reached statistical significance (**Figure 5B**).

**Figure 5.**
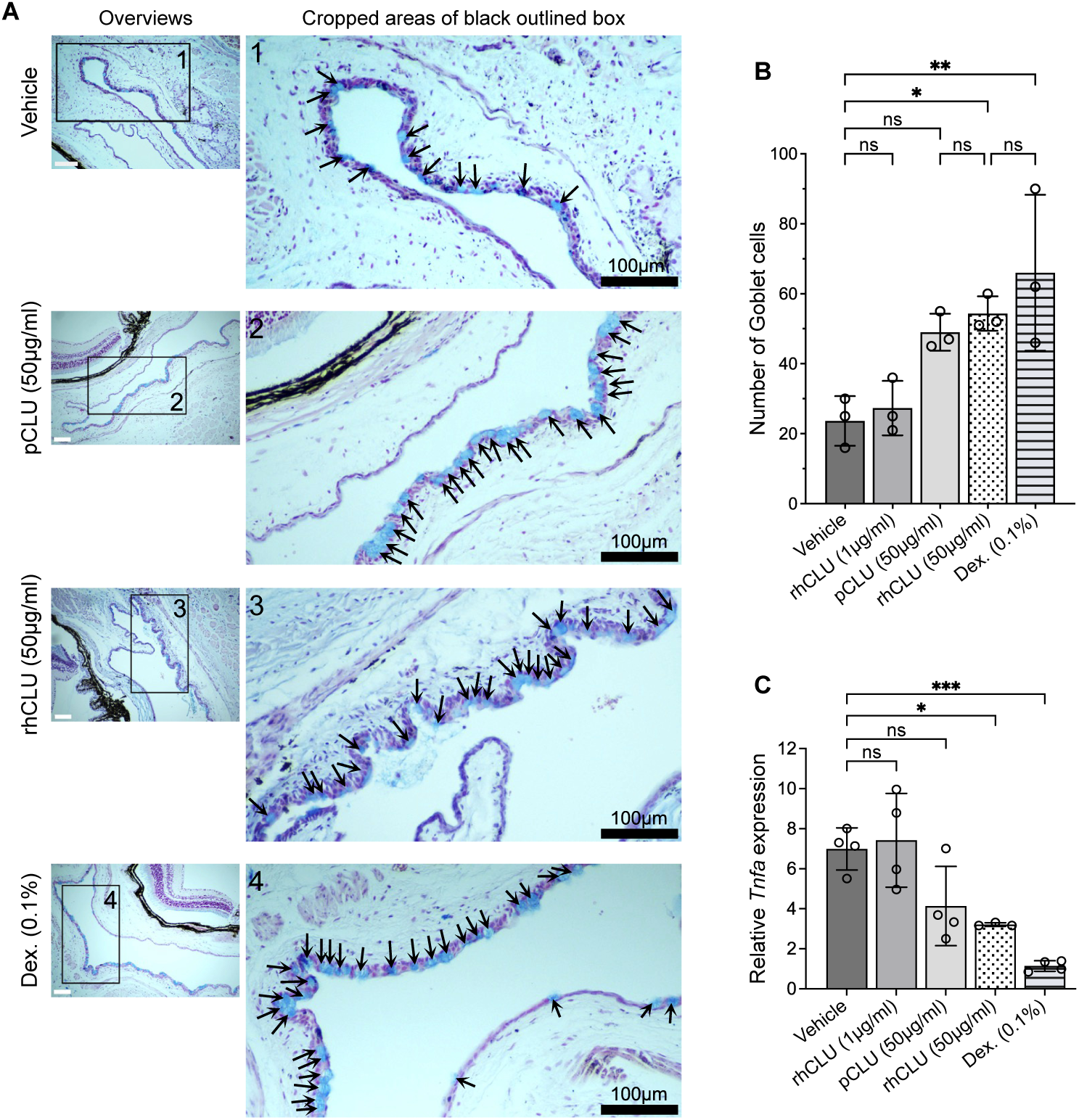
CLU treatment promotes regeneration of conjunctival Goblet cells, correlating with reduced conjunctival *Tnfα* expression levels in mice with Sjögreńs disease. (**A**) Representative alcian blue-stained conjuctiva tissue sections of *Thbs1*-deficient mice from groups with indicated treatments. Cropped images represent only marked areas in overviews of conjunctiva that are representative of each group. Blue-stained Goblet cells are indicated by arrows. (**B**) Goblet cell densities are represented as number of goblet cells counted in different treatment groups. (n= 3 mice/group). (**C**) Conjunctival *Tnfα* expression levels after three weeks of CLU treatment were determined from conjunctival RNA of indicated treatment groups using the SYBR Green real-time PCR method. Dexamethasone (Dex.) served as positive control with established anti-inflammatory effects. Error bars represent ± SEM (ns ≥ 0.05, *p < 0.05, **p < 0.01, ***p < 0.001).

We have previously reported that inflammatory cytokines induce conjunctival goblet cell apoptosis and contribute to their loss in *Thbs1*-deficient mice ^40,54^. To determine if increased goblet cell numbers correlate with reduced inflammation, we evaluated the expression of *Tnfa* as a marker of inflammation by RT-PCR, using RNA isolated from conjunctival tissues harvested from different treatment groups at the study endpoint. Consistent with its known anti-inflammatory effect, dexamethasone treatment resulted in significantly reduced expression of *Tnfa* as compared to vehicle treatment (**Figure 5C**). While treatment with 1 µg/ml of rhCLU showed no significant difference, treatment with 50 µg/ml of rhCLU resulted in significantly reduced *Tnfa* expression levels as compared to the vehicle control (**Figure 5C**). Together, these results highlight the potential of CLU to mitigate conjunctival inflammation.

### CLU treatment reduces corneal inflammation associated with Sjögreńs disease

Since ocular surface disease-related inflammation affects both cornea and conjunctiva, we investigated the effect of clusterin on corneal inflammation to gain a deeper understanding of its anti-inflammatory potential. Corneal inflammation in *Thbs1*-deficient mice is associated with an increased expression of *Tnfa* ^39,40,46,55^, and release of this cytokine from the surface of the cells as a mature protein is mediated by the action of the metalloproteinase ADAM17 (also known as TNFα-converting enzyme or TACE) ^56,57^. Recently, increased expression of ADAM17 was reported in corneal epithelium of *Thbs1*-deficient mice ^55^. To evaluate the effect of topically administered CLU on corneal inflammation we examined ADAM17 expression in the corneal epithelium of *Thbs1*-deficient mice from each treatment group. As shown in **Figure 6A**, ADAM17 immunostaining, indicated by the intense red fluorescence, was detected in the corneal epithelium of vehicle-treated mice, while the intensity of this staining was reduced in mice treated with CLU (50 µg/ml pCLU or rhCLU) or 0.1% dexamethasone (positive control).

**Figure 6.**
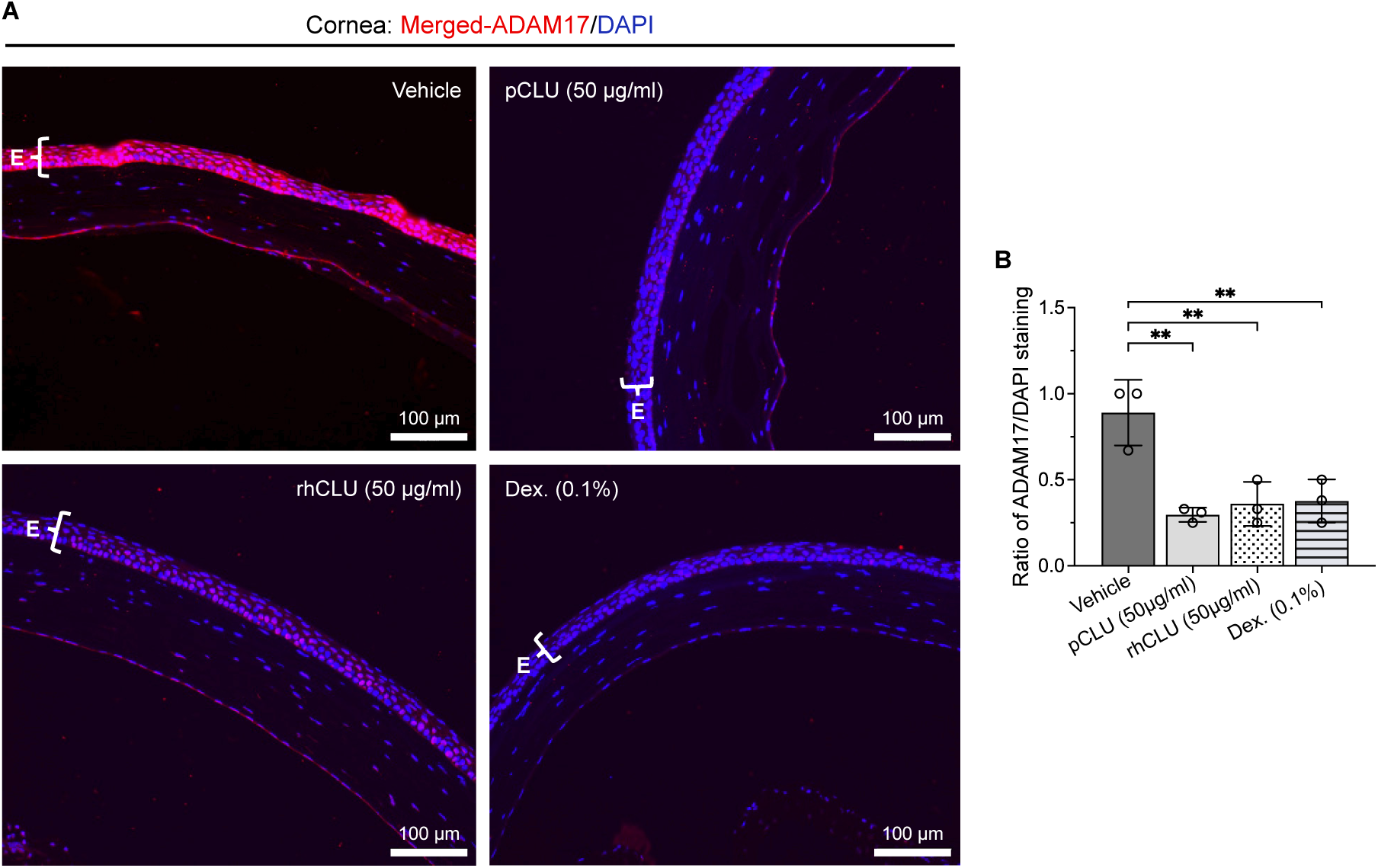
Topical CLU inhibits corneal ADAM17 expression levels in a mouse model of Sjögren’s disease. Eyes harvested from *Thbs1*-deficient mice treated with vehicle or 50µg clusterin (pCLU or rCLU) or 0.1 % dexamethasone (Dex.) (positive control) for up to 3 weeks (5 days/week) were harvested, and parafin-embedded sections were immunostained for ADAM17 as described in the methods. (**A**) Representative immunofluorescence images of ADAM17 expression (red) in the corneal Epithelium (E) from the indicated treatment groups are shown. (**B**) Quantitative analysis of ADAM17 staining within the corneal epithelium presented as the ratio of ADAM17 (red) to DAPI (blue) staining. Data presented as mean ± SEM (**p < 0.01).

Quantitative analysis (**Figure 6B**) confirmed a significant reduction in the amount of immunoreactive ADAM17 in the corneal epithelium of CLU (66% and 60% decline in pCLU and rhCLU-treated mice, respectively) and dexamethasone-treated mice (57%), as compared to vehicle-treated mice.

In summary, our results clearly demonstrate that, similar to topically administered dexamethasone, CLU (pCLU and rhCLU) effectively reduces ADAM17 immunoreactivity in the corneal epithelium and reduces *Tnfa* expression in the conjunctiva, supporting its anti-inflammatory effect in the treatment of chronic ocular surface disease.

## Discussion

Autoimmune SjD is associated with the development of chronic inflammatory dry eye disease, ranging from moderate to severe ^1,9^, which compromises the ocular surface, particularly affecting the corneal epithelium, as well as corneal nerves ^10,43,58^. Current approved therapeutic options to treat dry eye disease target inflammation with an aim to indirectly restore inflammation-induced epithelial and nerve damage. However, this approach has not proven effective in reversing corneal epitheliopathy ^59–61^. In fact, in one study, topical steroid treatment failed to improve corneal epitheliopathy in patients with reduced corneal nerve density, highlighting the inadequacy of targeting inflammation and the integral role of corneal nerves in maintaining the integrity of corneal epithelial barrier ^62^. Our study demonstrates that topical application of CLU has the potential to address challenges in treating chronic dry eye disease, as it not only helps seal corneal epithelial damage and reverse epitheliopathy but also supports regeneration of damaged corneal nerves while reducing ocular surface inflammation.

Several studies have demonstrated that the bidirectional interaction between the epithelium and nerves plays a crucial role in maintaining ocular surface health and visual function ^62–64^. This reciprocal interaction between corneal epithelium and nerves is further highlighted by the negative correlation observed between corneal epitheliopathy and corneal nerve density in patients with SjD ^10^. Furthermore, reduced corneal nerve density in SjD is also associated with increased density of inflammatory cells in the corneas ^43^. These studies indicate that more severe ocular surface damage is associated with lower nerve density. Consistent with these studies, our results demonstrate that the topical application of CLU in mice with SjD significantly improved corneal epithelial barrier integrity and increased subbasal corneal nerve density in both the peripheral and central areas. Moreover, as reported in SjD patients, we detected an inverse correlation between corneal epitheliopathy and corneal nerve density ^10^. A similar correlation was observed in patients with neurotrophic keratopathy treated with recombinant human nerve growth factor (rhNGF) ^50^. Although CLU has been shown to possess neuroprotective properties, no direct interactions between clusterin and nerve cells, nerve fibers or nerve function have been identified yet ^65–67^. In the cornea, CLU may indirectly influence nerve function by sealing the damaged epithelium, which allows the regenerated epithelium to release growth factors that promote nerve growth ^18,22^. Future studies will be important to uncover the precise mechanisms by which CLU affects corneal nerves ^68^.

Furthermore, our study revealed an interesting phenomenon: three weeks after CLU application, the improvement in corneal nerve density was significantly greater in the peripheral areas compared to the central areas. This improvement was even more pronounced in the CLU-treated group than in the steroid-treated group. The mouse model allowed us to precisely differentiate between the peripheral and central regions. In contrast, corneal nerve density data from human patients, typically collected using the non-invasive imaging technique called in vivo confocal microscopy (IVCM), mostly focuses on the central cornea ^50,62,69^. In contrast, a clinical study on corneal nerve growth following cenegermin (rhNGF) application in patients with neurotrophic keratopathy included IVCM data from both the central and peripheral cornea^70^. In this study, subbasal nerve density initially increased significantly in the periphery, while it only began increasing in the central cornea after several months of treatment ^70^. Our findings are consistent with the observed centripetal direction of corneal nerve regeneration in human patients ^70^. Taken together, these results suggest that corneal nerve regeneration occurs from the periphery toward the center. Whether the epithelium plays a distinct role in clusterin-mediated nerve regeneration in the periphery versus the center remains an intriguing and unanswered question. Known local differences in central and peripheral corneal cell layer thickness, epithelial regenerative capacity, and epithelial cell density may influence epithelial-nerve interactions and regeneration dynamics ^71,72^.

Studies have reported that sensory nerve-derived neuropeptide CGRP accelerates corneal epithelial growth and promotes wound healing ^51,73^. Furthermore, this neuropeptide was reported to exert an immunomodulatory effect, *in vivo*, by inhibiting antigen presentation by Langerhans cells in the skin ^74^. Consistently, corneal nerve-derived CGRP is now reported to play a cytoprotective and anti-inflammatory role in corneal wound healing ^17^. Our observation of a reduced proportion of CGRP-positive corneal nerves in *Thbs1*-deficient mice that develop ocular surface disease is, therefore, consistent with the wound-healing and anti-inflammatory roles of CGRP. The ability of topically applied CLU to increase the proportion of CGRP-positive corneal nerves not only aligns with its ability to support the healing of epithelial barrier damage but also suggests a role in countering ocular surface inflammation. This possibility is confirmed by the reversal of inflammation-mediated conjunctival goblet cell loss and reduced expression of the inflammatory marker *Tnfa* in the conjunctiva. In the cornea, epithelial damage is associated with an increased expression of a pro-inflammatory marker ADAM17 that amplifies inflammation by cleaving cell membrane TNFA ^75–77^. Increased expression of ADAM17 was reported in corneal epithelium of *Thbs1*-deficient mice, consistent with the increased expression of *Tnfa* ^39,46,55^. In our study, topically applied CLU also reduced corneal positivity for immunoreactive ADAM17, confirming the targeting of corneal inflammation. Inhibition of *Adam17* expression has shown a downstream effect of reducing MMP9 activity ^76,78,79^. Therefore, the observed *in vivo* anti-inflammatory effect of CLU in the cornea is consistent with its previously reported *in vitro* ability to inhibit the enzymatic activity of pro-inflammatory MMP9 ^31^. Overall, the anti-inflammatory effect of CLU in our study is consistent with the findings of an exacerbated inflammatory response in autoimmune disease models induced in mice lacking *Clu* ^80–82^, as well as other reports of its anti-inflammatory effects ^83,84^.

Chronic inflammation plays a central role in the development and persistence of ocular surface disease, and the inflammatory cascade can cause tissue damage and corneal nerve sensitization, effectively reducing pain threshold to perpetuate the dry eye disease cycle in the corneal and conjunctival tissues ^15,85^. As a result, currently approved treatments for dry eye primarily target inflammation. However, they offer limited efficacy in repairing corneal epithelial damage and providing symptomatic relief ^2,12^. In addition to sealing damaged corneal epithelium, our results clearly demonstrate an anti-inflammatory effect of CLU in both the cornea and conjunctiva. Thus, while CLU effectively seals acutely damaged corneal epithelium ^35,36^, our results establish that it also has a targeted sealing effect in chronic inflammatory conditions and improves corneal nerve density, potentially alleviating neurosensory impairments that contribute to symptoms. These effects, along with its anti-inflammatory impact on the ocular surface, are comparable to, or even surpass, those of steroid treatments. Thus, CLU could serve as a viable alternative to steroids or as a complementary treatment in managing ocular surface diseases. Based on its efficacy observed in this study, we propose CLU as a beneficial biologic for treating a wide range of ocular inflammatory conditions involving corneal epitheliopathy and nerve loss, including dry eye disease, laser refractive surgeries, erosions, trauma, corneal infections and neurotrophic keratopathy.

## Materials and methods

### Animals

Twelve-week-old, homozygous Thrombospondin-1 (*Thbs1*) deficient mice (B6.129S2-*Thbs1^tm1^*^Hyn^/J, cat. # 6141) and the corresponding wild-type (WT) mice (C57BL/6J, cat. # 664) were obtained from Jackson Laboratories (Bar Harbor, Maine, USA). All experimental procedures were conducted in accordance with the ARVO Statement for the Use of Animals in Ophthalmic and Vision Research and were approved by the Institutional Animal Care and Use Committee (protocol TR201900023) at Boston University School of Medicine (Boston, MA, USA). Throughout the study, animals were housed in a pathogen-free environment with veterinary care provided by the animal facility at Boston University School of Medicine. After reaching the experimental endpoint, mice were euthanized with CO_2_ gas according to the American Veterinary Medical Association Guidelines for the Euthanasia of Animals.

### Preclinical study design

The therapeutic efficacy of CLU in resolving chronic ocular surface inflammation was evaluated in a double-blinded study using male and female *Thbs1*-deficient mice. These mice spontaneously develop chronic ocular surface inflammation resembling SjD, which affects the lacrimal gland, cornea, and conjunctiva, with symptoms fully established by 12 weeks of age^42^. Consequently, 12-week-old *Thbs1*-deficient mice were selected for this preclinical study. A total of seven groups of 12-week-old *Thbs1*-deficient mice (n = 5 per group) were included. Five groups of male mice received topical treatments with 5 µl of the respective therapeutic eyedrop per eye, administered twice daily (b.i.d.) for five days per week over a three-week period. The treatment groups included: (1) vehicle (PBS), (2) rhCLU, 1 µg/ml, (3) pCLU, 50 µg/ml, (4) rhCLU, 50 µg/ml, and (5) 0.1% dexamethasone (Sigma-Aldrich, St. Louis, MO, USA), as a positive control. pCLU was purified from human plasma using immunoaffinity chromatography, as described by ^86^, while rhCLU was generated in our laboratory according to the method described ^87^. Corneal epithelial barrier integrity was assessed prior to initiating treatment (baseline) and monitored weekly throughout the study. At the study endpoint, mice were euthanized, and various tissues, including whole eyes with eyelids, corneas and conjunctiva, were harvested for further analyses.

### Corneal fluorescein staining

Corneal epithelial barrier integrity was performed using corneal fluorescein staining (CFS), as previously described ^39^. Briefly, 3 µl of 1% sodium fluorescein (Sigma-Aldrich, St. Louis, MO, USA) was applied to both eyes of isoflurane-anesthetized mice for 1 minute, followed by a brief wash to remove excess dye. The staining of the corneal epithelium was then evaluated using a slit-lamp microscope (Haag-Streit, Köniz, Switzerland) equipped with a cobalt blue light. Punctate fluorescein staining was graded weekly for both eyes of each mouse using the standardized National Eye Institute (NEI) grading system of 0-3 for each of the five areas of the cornea ^88^. Finally, CFS scores were normalized to the baseline mean of each treatment group for comparison.

### Immunohistochemistry

At the study endpoint, one eyeball from each mouse was enucleated and corneas were dissected and immediately fixed in 2% paraformaldehyde for 1 hour on ice. Radial incisions were made in fixed corneas to allow creating flat mounts. Fixation was followed by washes in PBS and permeabilization in PBS/1% Triton X-100 for 30 minutes and blocking with a solution of 1% BSA, 0.3% Triton X-100 and 10% donkey serum for 1 hour. Corneas were incubated with primary antibodies for beta-3-tubulin for 24 hours at 4°C or CGRP for 72 hours at 4°C prior to staining with anti-beta-3-tubulin antibody. Following washes with wash buffer corneas were incubated with fluorescence-conjugated secondary antibodies for 1 hour at room temperature (RT) each. After final washes, corneas were mounted in polyvinyl alcohol mounting medium (Sigma-Aldrich, St. Louis, MO, USA).

To visualize in situ ADAM17 immunoreactivity in the corneal epithelium, eyes harvested at the end of the study period were fixed in 4% paraformaldehyde before paraffin embedding. Sections 5 µm in thickness (n = 3 per group) were used for immunostaining. Sections were blocked at room temperature (RT) for 30 minutes with 10% goat serum in PBS, followed by 2% bovine serum albumin (BSA) (Sigma-Aldrich, St. Louis, MO, USA), 0.1% Triton-X100 in PBS for 1 hour (h), and incubated overnight at 4°C with primary anti-ADAM17 antibody in PBS-BSA. Tissues were washed with PBS-0.05% Tween-20 and incubated for 1 hour at RT with fluorescence-conjugated secondary antibody. Tissues were further washed with PBS-0.05% Tween-20 and counterstained with DAPI.

Primary antibodies were rabbit anti-TuJ1 (1:500; Cat. # T2200, Sigma-Aldrich, St. Louis, MO, USA), mouse anti-CGRP (1:200; Cat. # sc-57053, Santa Cruz Biotechnology, Dallas, USA) and rabbit anti-ADAM17 (1:50; Cat. # sc-13973, Santa-Cruz Biotechnology, Dallas, USA). Secondary antibodies included Alexa Fluor-488-conjugated donkey anti-rabbit IgG (1:1000; Cat.# A-21206, Invitrogen, Carlsbad, CA, USA), DyLight-650-conjugated donkey anti-mouse IgG (1:200; Cat.# ab98769, Abcam, Cambridge, UK) and Alexa Fluor-568-conjugated goat anti-rabbit IgG (1;500, Cat.# A-11011, Invitrogen, Carlsbad, CA, USA).

### Microscopy and image analysis

To visualize immunostained corneal nerves, Z-stacks of images were acquired using a Zeiss LSM 710-Live Duo Confocal microscope (Carl Zeiss, Oberkochen, Germany). Overview images of corneal nerve morphology in the peripheral cornea were acquired with a Zeiss Plan-Apochromat 10x/0,45 M27 (0.6 zoom) objective (Carl Zeiss, Oberkochen, Germany) and corneal nerves in selected central and peripheral regions were imaged using a Plan-Apochromat 20x/0,8 Ph2 M27 (Carl Zeiss, Oberkochen, Germany) objective. The same acquisition settings were consistently applied to all samples within the same biological replicate. Image processing, including brightness and contrast, were adjusted with Fiji ImageJ ^89^. Corneal nerve density was quantified based on beta-3-tubulin immunostaining of wholemount corneas. For analysis, each cornea was divided into four central and four peripheral regions as shown in figure 2A. The central region, characterized by a whorl-like nerve structure, was imaged using four adjacent Z-stack scans (425 µm x 425 µm each). Additionally, one Z-stack scan was randomly taken from each of the four peripheral regions. In total, 25 images from both the central and peripheral corneal regions were analyzed per treatment group. Maximum intensity projections of the sub-basal corneal nerve zone were then used for nerve density measurements and nerves were segmented using the ‘Ridge detection’ plugin in Fiji ImageJ ^90^. Consistent segmentation settings were applied across all treatment groups to ensure accurate comparisons. Finally, nerve density was calculated as the percentage of the area covered by nerves relative to the total image area. Quantification of CGRP-staining was performed in corneas immunostained for beta-3-tubulin and CGRP. First, the total length of beta-3-tubulin positive sub-basal corneal nerves was measured using the ImageJ ‘Ridge detection’ tool ^90^. Second, the length of CGRP-positive nerves was manually traced using the Fiji ImageJ ‘NeuronJ’ plugin ^91^. The percentage of CGRP-positive nerve fibers was calculated relative to the total beta-3-tubulin-positive nerve length in each image.

Fluorescence staining of ADAM17 in corneal epithelium was evaluated using a fluorescence microscope (Nikon Eclipse E800, Nikon, Japan) and images were captured with a MicroPublisher 6 camera. Images were further analyzed using Fiji ImageJ software ^89^ to determine relative ADAM17 staining (red) to nuclear DAPI (blue) staining. Fluorescence intensity values for both colors within the same ROI were used to calculate the ratio.

### Goblet cell density

At the end of the 3-week study period, one whole eye with lid was harvested from three mice per treatment group, fixed in 4% paraformaldehyde and embedded in paraffin. Sagittal sections (5 µm) from the center of the eye were cut, stained with alcian blue/PAS solution (Vector Laboratories, CA, USA) and mounted in Permount medium. Representative sections of the bulbar and palpebral conjunctiva, showing filled goblet cells stained blue, were captured using a Plan Fluor 10x/0.30 DIC WD objective (Nikon, Tokyo, Japan). The mean goblet cell number in both bulbar and palpebral conjunctiva were determined for two sections per eyeball.

### Real-time PCR

At the end of the study period (3 weeks), conjunctiva collected from mice in each treatment group was pooled and total RNA was isolated using TRIzol Reagent. RNA concentration and quality were assessed with a NanoDrop device. cDNA synthesis was performed using the SuperScript VILO cDNA synthesis kit (Thermo Fisher Scientific, Waltham, USA) according to the manufacturer’s instructions. 1 µg of RNA per group was subjected to reverse transcription under the following thermal conditions: 25°C for 10 minutes, 50°C for 10 minutes, and 85°C for 5 minutes. For amplification of *Tnfα* and *Gapdh* gene transcripts, a semi-quantitative PCR was conducted using diluted cDNA (1:10), gene-specific primers (*Tnfα*: F-5’-GGCCTCCCTC TCATCAGTTCTATG-3’, R-5’-GTTTGCTACGACGTGGGCTACA-3’; *Gapdh*: F-5’-CGAG AATGGGAAGCTTGTCA-3’, R-5’-AGACACCAGTAGACTCCACGACAT-3’) and the SYBR Green PCR Master Mix (Thermo Fisher Scientific, Waltham, USA) with the following thermal profile: 3 minutes at 95°C, followed by 40 cycles of 20 seconds at 95°C, 30 seconds at 55°C, and 40 seconds at 72°C. Fluorescence signals were recorded during each cycle and analyzed using StepOne Software v2.3 (Thermo Fisher Scientific, Waltham, USA). Melting curve analysis was performed to confirm the specificity of each RT-PCR reaction. Threshold cycle values were used to evaluate gene expression levels relative to the reference gene Gapdh.

### Statistics

Statistical analyses were performed by using the GraphPad PRISM software (GraphPad Software, version 10, La Jolla, USA). Normal distribution of the data was assessed using D’Agostino and Pearson test. For comparisons between two groups with normally distributed data, an unpaired two-tailed Student’s t-test was performed. When data did not follow Gaussian distribution, the Mann-Whitney test was applied. For comparisons between more than two groups, a one-way ANOVA test was applied. Pearson’s correlation coefficient was used to evaluate correlation between corneal fluorescein staining score and corneal nerve density. Error bars show ± SEM and differences were considered significant when p < 0.05 (*p < 0.05; **p < 0.01; ***p < 0.001; ****p < 0.0001).

## Acknowledgments

Boston University received a subcontract from NIH grant R41EY034396 awarded to Proteris Biotech.

## Author contributions

Conceptualization and experimental design: M.E.F., S.M., Funding Acquisition: M.E.F., S.M., Experimental set up, data acquisition and analysis: J.F., T.F.N, S.G., Project Administration: M.E.F., S.M., Resources: M.R.W., Supervision: M.E.F., S.M., Writing – Original Draft Preparation: J.F., Review & Editing: J.F., T.F.N, S.G., M.E.F., S.M.

## Data availability

All data generated or analyzed during this study are included in this published article.

## Competing interests

M.E.F. is a co-founder of Proteris Biotech and an employee of the company, serving as Chief Scientific Officer. S.M. and M.R.W. hold equity in Proteris Biotech and serve on the Medical and Scientific Advisory Board. M.E.F. is named as a co-inventor on U.S. patent number 9241974 entitled “Clusterin Pharmaceuticals and Treatment Methods Using the Same” granted to the University of Southern California and optioned for exclusive license to Proteris Biotech. S.M. and M.E.F. are named as co-inventors on a preliminary U.S. patent application co-submitted by Boston University and Proteris Biotech, claiming some of the new findings of this study (63/734,859). All other authors declare that they do not have any competing interest.

## Additional information

Correspondence and requests for materials should be addressed to S.M. or M.E.F.

---

**Supplementary Figure 1.**
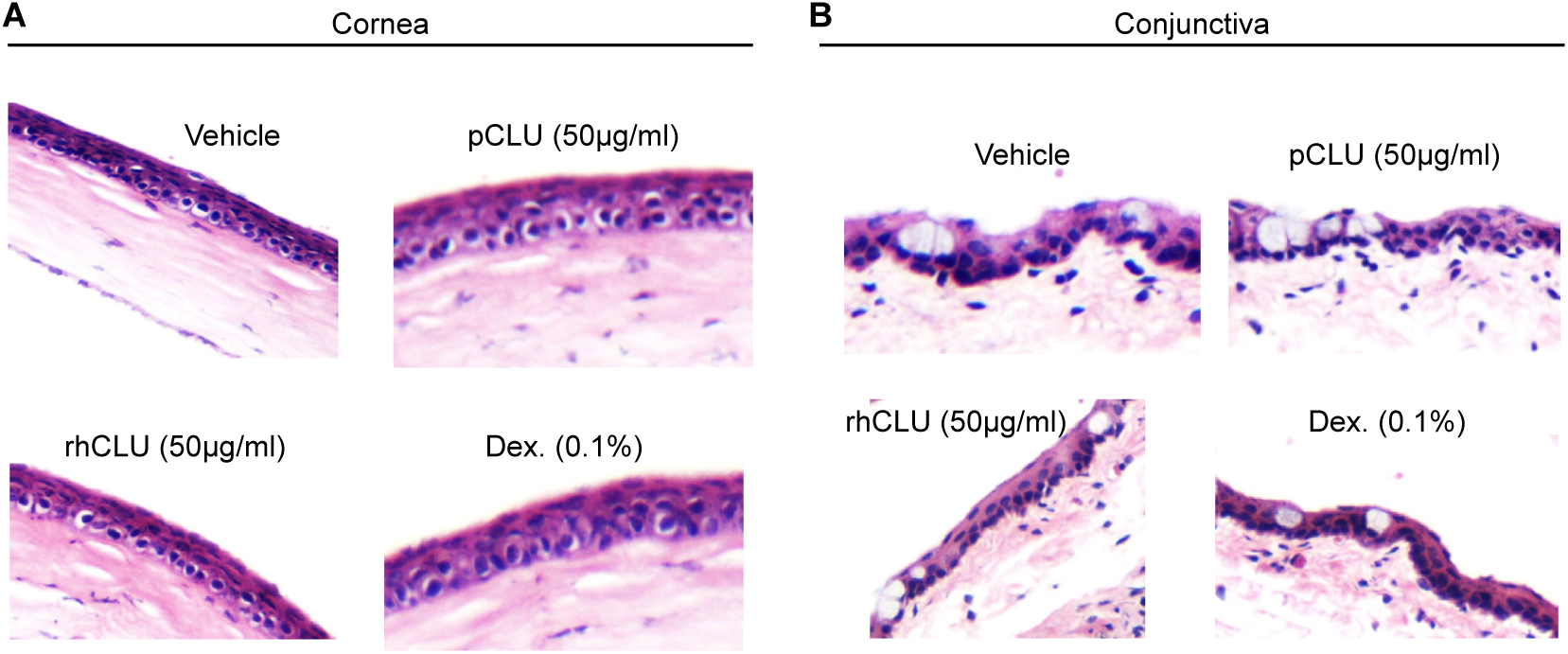
Corneal and conjunctival tissue morphology in Hematoxylin and Eosin (H&E) staining remains unchanged after topical application of CLU. Representative H&E images (20x magnification) of (**A**) corneal and (**B**) conjunctival tissues from indicated treatment groups showed no changes in tissue morphology and no inflammatory infiltrates.

